# A diet enriched in highly poly-unsaturated fatty acids alters immune cell lipid composition and ferroptosis susceptibility

**DOI:** 10.64898/2026.10.20.760583

**Authors:** Pooranee K. Morgan, Yangsong Xu, Thomas J. C. Collins, Camilla Bertuzzo Veiga, Natalie Mellett, Kevin Huynh, Peter J. Meikle, Andrew J. Murphy, Graeme I. Lancaster

## Abstract

Ferroptosis is a regulated form of cell death that is dependent upon the peroxidation of phospholipids containing poly-unsaturated fatty acids (PL-PUFAs), particularly those with ≥4 double bonds. PL-PUFA content is dramatically different between immune cell types which directly influences their sensitivity to ferroptosis. Modulating the PL-PUFA content of various cells is becoming an attractive approach to increase the efficacy of therapies designed to target ferroptosis. In this study we aimed to determine whether a diet enriched in highly unsaturated PUFAs alters immune cell lipid composition and, consequently, the susceptibility to ferroptosis. To address this, mice were fed either a diet enriched in eicosapentaenoic acid (EPA) and docosahexaenoic acid (DHA) (High PUFA diet) or a standard control diet (Control diet) for eight weeks. Immune cells were then isolated from the bone marrow, blood, and lymph nodes for lipidomic analysis and sensitivity to ferroptosis. The High PUFA diet resulted in the widespread incorporation of EPA and DHA into multiple PL species across all immune cell types. However, the magnitude of incorporation differed between cell populations, with B cells exhibiting the greatest enrichment in PL-PUFA-EPA and PL-PUFA-DHA species, followed by monocytes and neutrophils, while T cells showed the lowest degree of enrichment. Importantly, these compositional changes were associated with increased susceptibility to ferroptosis in a cell and location specific manner. Collectively, these findings demonstrate that a diet enriched in highly unsaturated PUFAs increases PL-PUFAs in immune cells and broadly enhances immune cell susceptibility to ferroptosis. Therefore, altering dietary fatty acid composition may represent a strategy to influence ferroptosis sensitivity in conditions where either inducing or inhibiting ferroptosis is therapeutically desirable.

## Introduction

Ferroptosis is a form of regulated cell death executed due to an imbalance in anti-oxidant defense mechanisms and pro-oxidant processes causing the peroxidation of PL-PUFAs in the cell membrane resulting in an unrepairable breach in the cell membrane (1, 2, 3, 4). Because of its involvement in a wide variety of diseases, targeting ferroptosis therapeutically has become the subject of intense focus and therefore there is significant interest in determining ways to modulate the susceptibility of cells to ferroptosis depending on disease context. Triggering ferroptosis is being investigated to treat cancer (5), while inhibiting ferroptosis may be relevant in ischemia reperfusion injury and neurodegeneration (6). While the pathways that control ferroptosis are multifactorial, PL-PUFAs, as the executioners of ferroptosis, play a primary role (1, 2, 4). Accordingly, altering PL-PUFA levels profoundly impacts the cellular sensitivity to ferroptosis (2, 4, 7).

The abundance of highly unsaturated PL-PUFAs, in particular arachidonic acid (AA; 20:4 n6) and docosahexaenoic acid (DHA; 22:6 n3), is dramatically different between immune cell subsets (7). T cells and B cells have markedly higher levels of PL-PUFA-AA and PL-PUFA-DHA compared to myeloid cells, especially neutrophils and eosinophils (7). Notably, neutrophils, which contain very low levels of PL-PUFA-AA and PL-PUFA-DHA, are almost entirely resistant to ferroptosis. These findings demonstrate a cell-intrinsic regulation of ferroptosis susceptibility based on their PL-PUFA content. While different cell types intrinsically regulate their PL-PUFA composition to achieve specific functions, we and others have shown that supplementing cells *in vitro* with mono-unsaturated fatty acids biases membrane PL composition away from PL-PUFAs and towards PL-MUFA and thereby reduces ferroptosis susceptibility, particularly in cells with a high basal PL-PUFA content (i.e. lymphoid cells) (7, 8). In contrast, supplementing cells *in vitro* with PUFAs, especially arachidonic acid (AA; 20:4 n6) and docosahexaenoic acid (DHA; 22:6 n3), sensitizes cells to ferroptosis, particularly those cells with low basal levels of PL-PUFAs (e.g. neutrophils) (4, 7, 9). Importantly, feeding mice a diet enriched in omega-3 fatty acids promotes ferroptosis in models of cancer and Crohn’s disease, suggesting that the PL-PUFA content of tissues can be altered by changes in diet (10, 11). How altering the fatty acid content of the diet influences immune cell sensitivity to ferroptosis is less well understood. Thus, we aimed to determine if a diet enriched in highly unsaturated PUFAs could increase immune cell PL-PUFA content and thereby increase their susceptibility to ferroptosis.

## Results

### Altering the PUFA content of macronutrient matched diets has no impact on overall metabolic profile

When supplemented to cells and animals, highly unsaturated PUFAs are especially well incorporated into the cellular lipidome (12); numerous studies have shown that diets enriched in marine omega-3 fatty acids such EPA and DHA increase EPA and DHA levels within immune cells (13). However, several important questions remain relatively unresolved: (i) are different types of immune cells differentially responsive to diet-induced alterations in lipid composition?; (ii) does immune cell location impact the response to diet-induced alterations in lipid composition?; and (iii) what are the precise lipidomic changes that occur in immune cells in response to a diet enriched in highly-unsaturated PUFAs. Therefore, to address these questions, we fed mice diets that were matched for overall fat content but contained either canola oil (enriched in the mono-unsaturated fatty acid oleic acid (18:1) and the PUFAs linoleic acid (18:2 n3) and α-linolenic acid (18:3 n6) (Control diet) or tuna oil (enriched in the highly unsaturated PUFAs EPA and DHA) (High PUFA diet) for 8 weeks (Fig. 1A and Table S1), after which immune cell lipid composition and ferroptosis susceptibility was determined (Fig. 1B). Mice gained the same amount of weight in response to either Control diet or High PUFA diet and no differences in circulating or bone marrow immune cell numbers were observed (Fig. 1C-E).

**Fig. 1.**
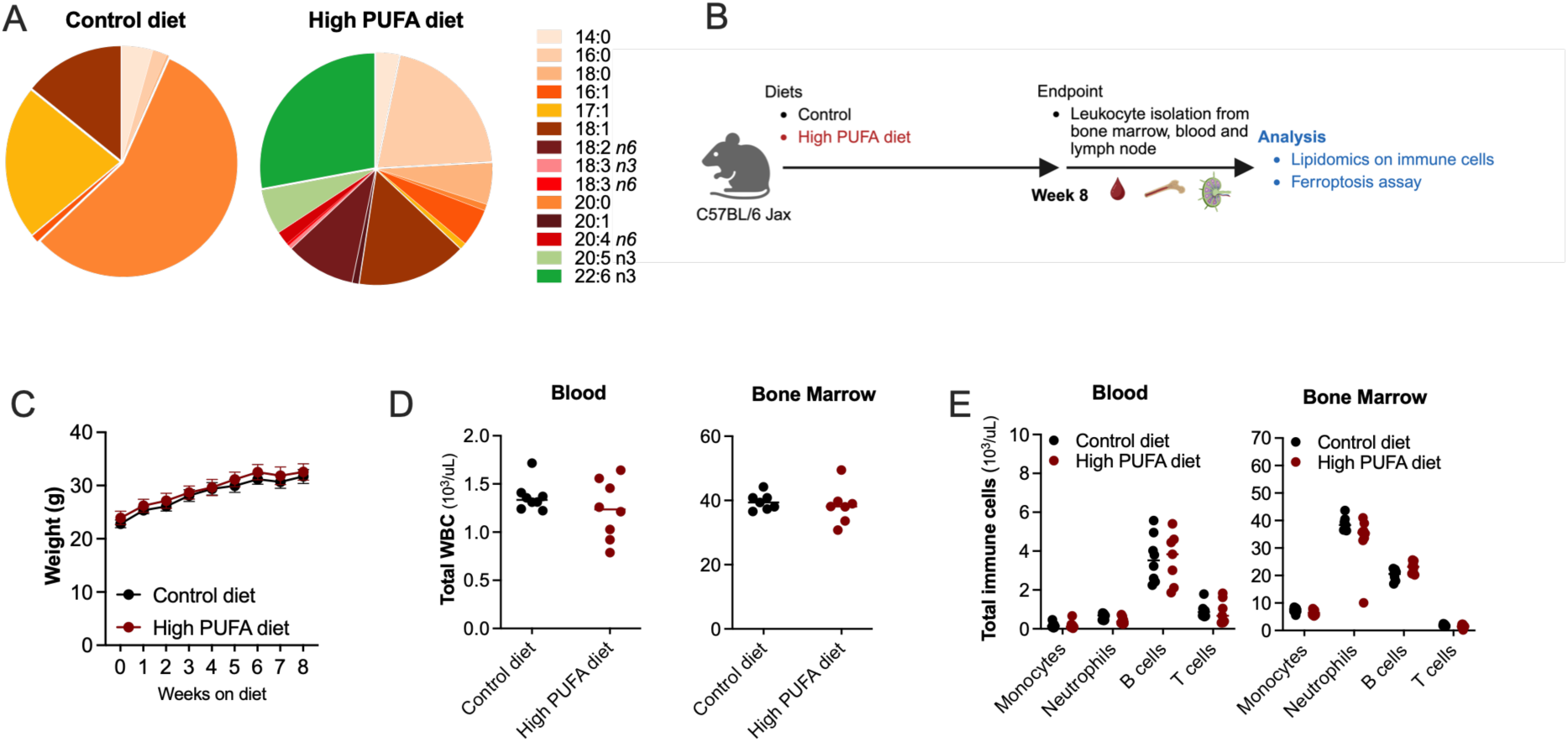
Study overview and characterisation of body weight and immune cell changes. **A** Fatty acid composition of the Control and High PUFA diets. **B** Schematic of the experimental workflow; C57Bl/6 mice were fed Control or High PUFA diets for 8 weeks. **C** Total body weight over time. **D** Total white blood cell count (WBC) in the blood and bone marrow post-8 weeks of diet. **E** Immune cells in the blood and bone marrow 8 weeks post diet. **C** n = 6 mice per group and **C-F** n = 7-8 mice per group.

### A diet enriched in highly unsaturated PUFAs dramatically alters immune cell lipidomes

To explore if the High PUFA diet altered the lipidomes of immune cells, T cells, B cells, neutrophils, and monocytes were isolated from blood and bone marrow, as well as total immune cells from lymph nodes, and their lipid compositions were quantified. To provide an overview of how the High PUFA diet altered the global lipidome of immune cells, differences were visualized on a UMAP (Fig. 2A-C). Consistent with our previous work, B cells, T cells, monocytes and neutrophils from bone marrow (Fig. 2A) and blood (Fig. 2B) clustered distinctly. Immune cells from bone marrow and lymph nodes displayed clear diet-dependent clustering (Fig. 2A and 2C). Interestingly, while there was still a degree of diet-dependent clustering between the immune cells from the blood, this was less dramatic and some overlap was observed (Fig. 2B). These observations indicate that the High PUFA diet profoundly alters immune cell lipid composition and that location (e.g. bone marrow, blood) may affect the extent of these diet-induced changes.

**Fig. 2.**
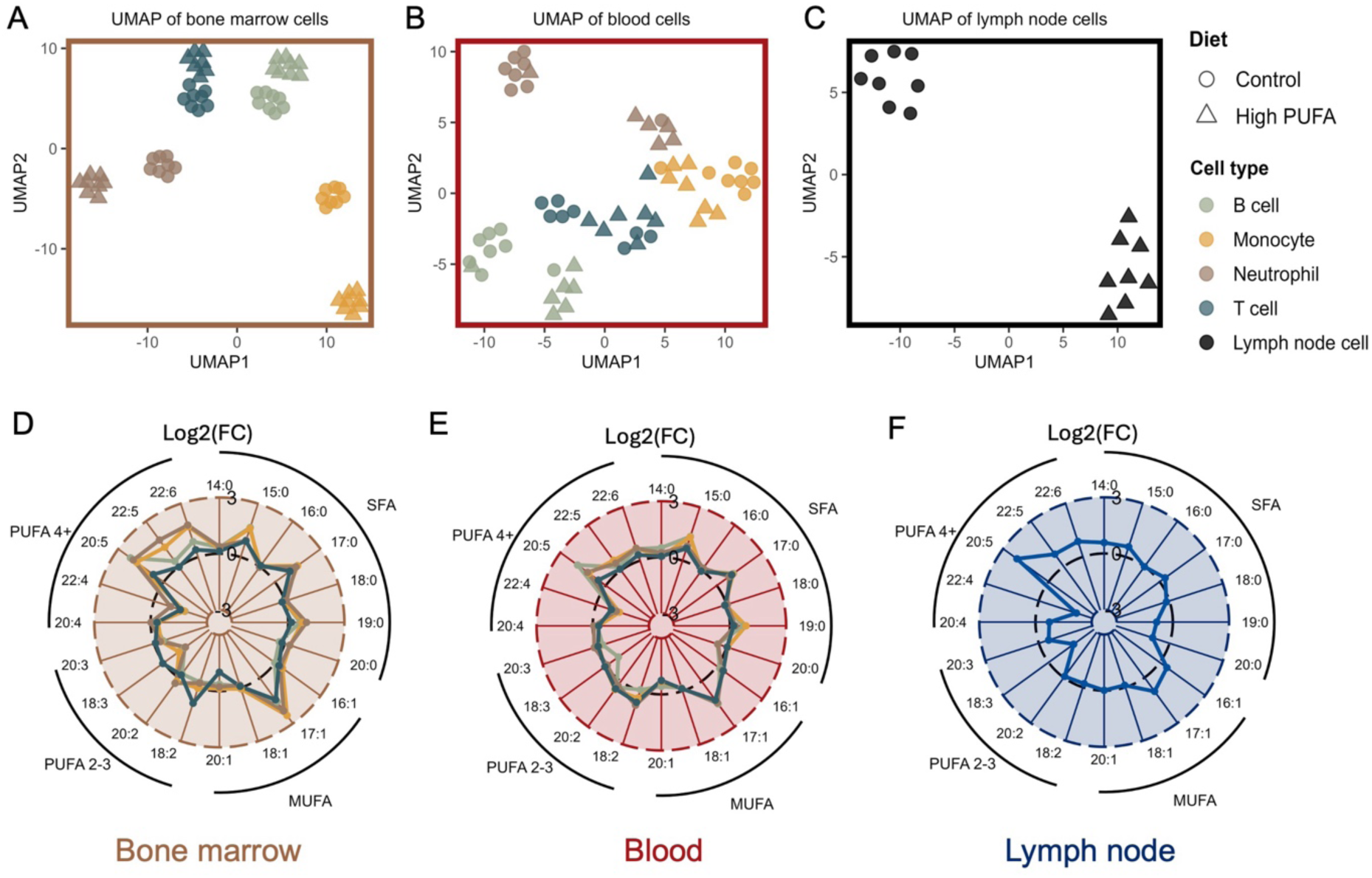
A High PUFA diet alters immune cell lipid composition. Isolated immune cells were subjected to mass spectrometry lipidomics. **A-C** UMAPs of the global lipidome of immune cells from the **A** bone marrow, **B** blood and **C** lymph nodes. **D-F** Radar plots showing changes in overall PL acyl chain composition in immune cells from the **D** bone marrow, **E** blood and **F** lymph nodes. In the radar plots, the effect of the High PUFA diet (coloured lines) is shown relative to the Control diet (black dotted line). **A** n = 7-8 mice per group; **B** n = 6-8 mice per group; **C** n = 8 mice per group.

Of the approximately 700 lipid species measured, several hundred were significantly different between the Control diet and High PUFA diet in immune cells from bone marrow, blood and lymph nodes (Fig. S1 to S3). Because the High PUFA diet is enriched in 20:5 and 22:6 PUFAs, we hypothesized that these fatty acids would be incorporated into cellular PLs, leading to an increase in PLs containing 20:5 and 22:6. As expected, PL acyl composition was markedly affected by the High PUFA diet, in particular PLs containing 20:5, 22:5, and 22:6 PUFAs were increased in all cell types from all locations (Fig. 2D-F and Fig. S4). These overall changes are the result of the incorporation of 20:5 and 22:6 into PLs from multiple classes, e.g. PC, PE, PE(P), and PI, highlighting the widespread incorporation of these dietary fatty acids into cellular PLs (Fig. 3A, B and Fig. S5). One of the most notable decreases in response to the High PUFA diet was PUFAs containing 3-4 C-C double bonds, in particular adrenic acid (22:4), with multiple 22:4-contining PL species being affected across all cell types from all locations (Fig. 2D-F and Fig. 3C). While the Control diet contained ∼4-fold higher levels of the SFA palmitic acid (16:0) and the MUFA oleic acid (18:1) compared with the High PUFA diet, the overall proportion of PLs containing 16:0 or 18:1 was similar between cell types (Fig. 2D-F).

**Fig. 3.**
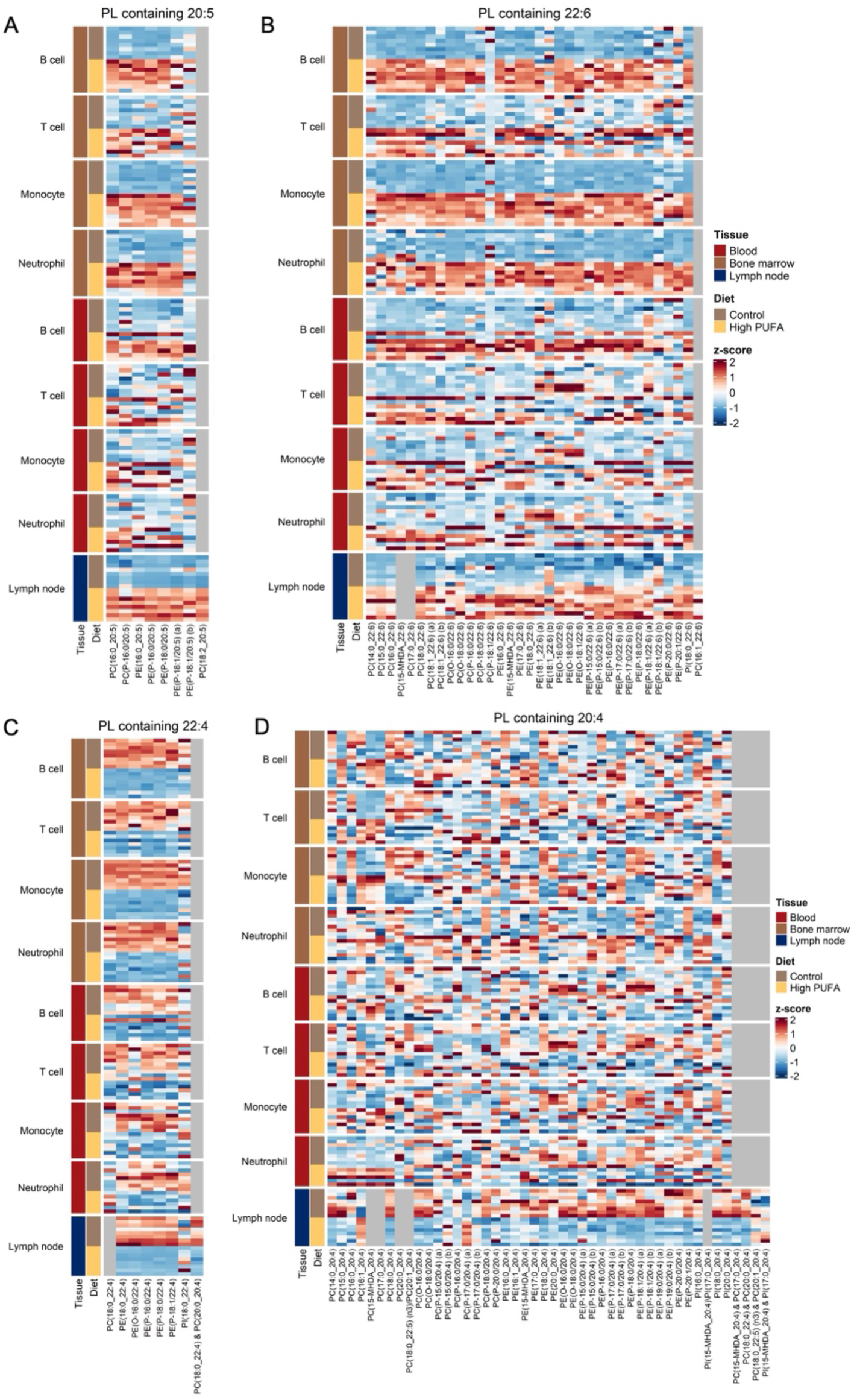
Effects of High PUFA diet on PLs species containing highly unsaturated PUFAs. **A-D** Heat maps showing changes in PL species containing **A** 20:5, **B** 22:6, **C** 22:4 and **D** 20:4 in immune cells from bone marrow, blood, and lymph nodes. For bone marrow immune cells, n = 7-8 mice per group; for blood immune cells, n = 6-8 mice per group; for lymph node immune cells, n = 8 mice per group. Data are presented as z-scores.

### High PUFA diet imparts cell- and tissue-specific lipid remodeling

While there is considerable commonality in the High PUFA diet-induced changes between different immune cell types and between locations (bone marrow, blood, lymph node), interestingly, there are also numerous cell-specific and tissue-specific changes. Firstly, the different immune cell types were differentially responsive to High PUFA diet-induced changes in lipid composition. This was particularly notable for T cells, with numerous individual 20:5- and 22:6-containing PL species (Fig. 3A, B and Fig. S5) and, consequently, total 20:5 and 22:6 levels (Fig. 2D-E and Fig. S4), being markedly less affected by the High PUFA diet in T cells compared with B cells, neutrophils, and monocytes. Secondly, and somewhat unexpectedly, immune cell location (i.e. blood *vs* bone marrow *vs* lymph node) had a marked effect on the degree of lipid re-modelling following the High PUFA diet. Specifically, changes in 20:5 and 22:6-containing PLs were generally more pronounced in immune cells from the bone marrow compared with the blood (Fig. 3A, B and Fig. S5). For example, 22:6-containing PLs in monocytes and neutrophils located in the bone marrow were increased ∼300% (Fig. S4A, E) but were not significantly increased monocytes and neutrophils in the blood (Fig. S4C, G). Finally, it is intriguing that PLs containing 20:4 were variably affected by the High PUFA diet in all cell types from bone marrow and blood (Fig. 3D). While in contrast, the majority of 20:4-containing PL species and overall 20:4 content were markedly lower following the High PUFA diet (Fig. 2F and Fig. 3D) in the lymph node cells. Collectively, this experiment demonstrates that immune cell lipid composition can be dynamically regulated by altering the composition of dietary fatty acids. Importantly, increasing the abundance of highly unsaturated PUFAs, i.e. EPA and DHA, in the diet has a widespread impact on immune cell PL acyl chain composition.

### High PUFA diet results in increased susceptibility of immune cells to lipid peroxidation and ferroptosis

Having shown that a High PUFA diet increased the abundance of PL-PUFA-EPA and PL-PUFA-DHA in various immune cell types, we next wanted to determine if these changes impacted the susceptibility to lipid peroxidation and ferroptosis. To this end, we treated immune cells from blood and bone marrow with the glutathione peroxidase 4 (GPX4) inhibitor ML210 and monitored lipid peroxidation (liperfluo fluorescence) and ferroptotic cell death. Consistent with our previous observations in immune cells obtained from mice fed a standard laboratory mouse diet (i.e. a diet that does not contain highly unsaturated PUFAs) (7), *ex vivo* treatment with ML210 for 6 hours triggered lipid peroxidation and ferroptosis in T cells but not in other immune cell types in mice fed the Control diet (Fig. 4). In line with our hypothesis, the High PUFA diet sensitized several immune cell types to lipid peroxidation and ferroptosis; however, the magnitude of this effect was cell-type specific (Fig. 4). Notably, B cells were the most affected, with B cells obtained from both the bone marrow and blood of mice fed the High PUFA diet being highly sensitized to lipid peroxidation and ferroptosis when GPX4 was inhibited (Fig. 4A, B). T cells from mice fed the High PUFA diet across the bone marrow, blood and lymph nodes were also significantly more susceptible to lipid peroxidation and ferroptosis (Fig. 4C, D, I), although the magnitude of this effect was not as marked as in B cells.

**Fig. 4.**
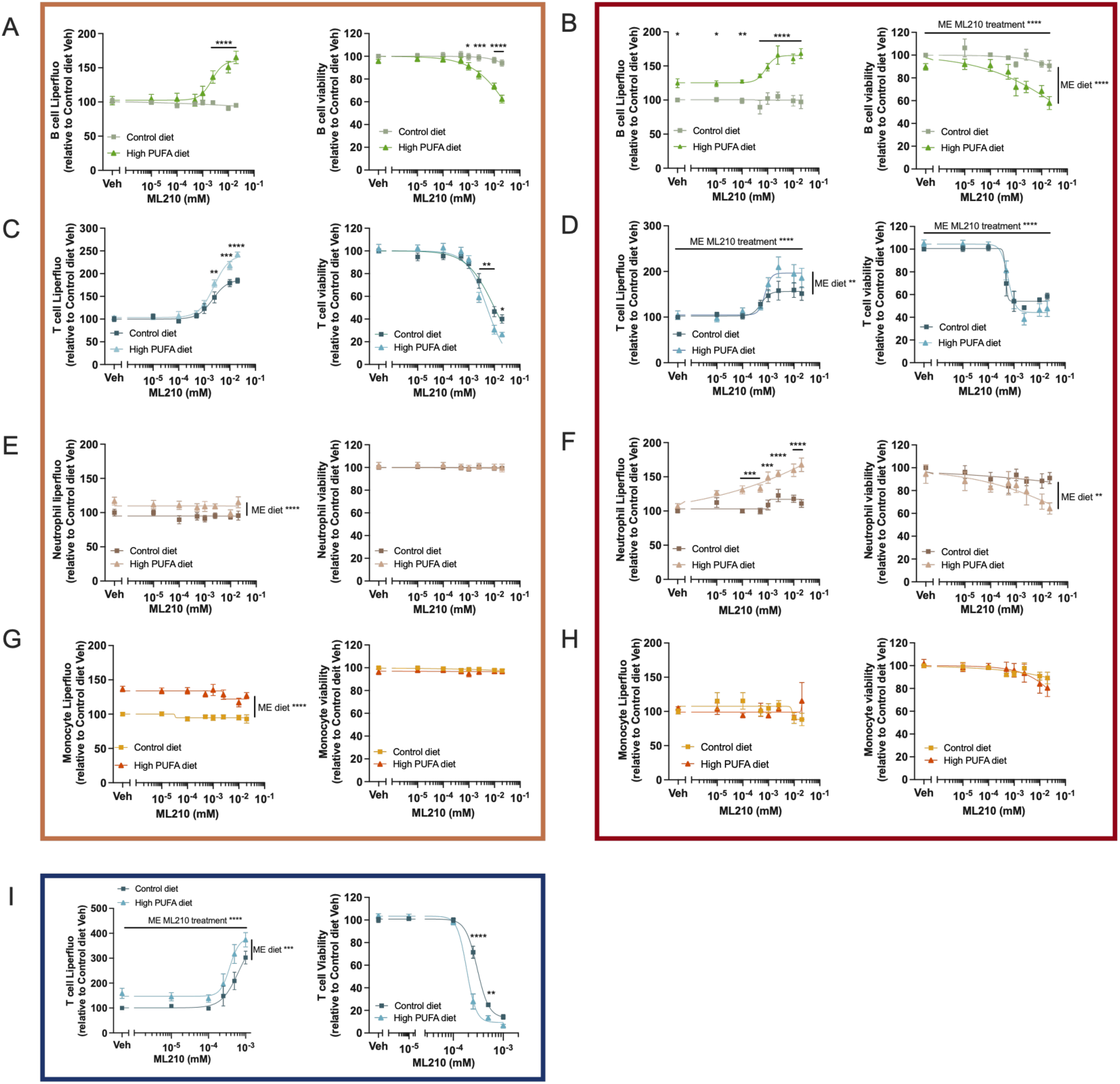
High PUFA diet sensitizes immune cells to lipid peroxidation and ferroptosis sensitivity. **A-I** Flow cytometry was used to quantify **Left panels**, lipid peroxidation (Liperfluo fluorescence) and **Right panels**, cell viability following *ex vivo* ML210 treatment for 6 h in **A, B** B cells, **C, D, I** T cells, **E, F** neutrophils and **G, H** monocytes isolated from the **A, C, E, G** bone marrow, **B, D, F, H** blood and **I** lymph nodes of mice fed either a Control diet or High PUFA diet. Statistical analysis was performed using two-way ANOVA with Tukey’s HSD test. *p<0.05, **p<0.01, ***p<0.001 and ****p < 0.0001 indicate significant pairwise comparisons between dietary conditions at specific ML210 doses following a significant interaction effect or, where specified, significant main effects (ME) for diet and ML210 treatment. n = 6 mice per group and data represent the mean ± SEM.

We have previously shown that monocytes, and in particular neutrophils, are relatively resistant to ferroptosis. Interestingly, following the High PUFA diet, neutrophils from the bone marrow displayed a GPX4 independent increase in lipid peroxidation, although this did not translate into increased ferroptotic cell death. Notably, neutrophils from the blood showed an increased susceptibility to lipid peroxidation and ferroptosis in response to GPX4 inhibition (Fig. 4E, F). No impact of the High PUFA diet was observed on GPX4 inhibited monocyte lipid peroxidation or ferroptosis in the bone marrow or blood, but similar to bone marrow neutrophils there was a GPX4 independent increase in lipid peroxidation in bone marrow monocytes (Fig. 4G, H). Collectively, these results show that feeding mice a diet enriched in highly unsaturated PUFAs is an effective strategy to alter the lipid composition and ferroptosis susceptibility of immune cells.

## Discussion

Modulating the pathways that control cell death has long been exploited to intervene in a plethora of life-threating and debilitating diseases. Ferroptosis, a form of cell death driven by lipid peroxidation of cell membrane PL-PUFAs, has become an attractive target to modulate depending on disease context (1, 3). The majority of previous studies that have sought to exploit ferroptosis as a means to eliminate unwanted cells have focused on inhibiting components of the pathways that prevent ferroptosis (e.g. GPX4, FSP1). Considering that membrane PL-PUFAs are well-established as the rate-limiting executioners of ferroptosis, it is surprising that so few studies have examined the potential of increasing the abundance of PL-PUFAs as means to enhance ferroptosis susceptibility. In this study we show that feeding mice a high PUFA diet increases the abundance of highly unsaturated PUFAs into the PL pool of immune cells, particularly lymphoid cells, rendering them susceptible to ferroptosis upon GPX4 inhibition.

The importance of highly unsaturated PUFAs in influencing the susceptibility of cells to ferroptosis is well established. This was formally shown by deleting enzymes that regulate the incorporation of fatty acids into PLs (2, 14) or by adding exogenous fatty acids to cells *in vitro* (4, 7, 8, 9). Studies are emerging that reveal diets can be modified to alter ferroptosis, whether it be to promote ferroptosis in solid tumors or limit ferroptosis in auto-inflammatory conditions (10, 11, 15). Wang et al, have shown that modulating the ratio of dietary PUFAs to MUFAs impacts T cell ferroptosis, which has several consequences on immune responses (16). These studies provide evidence that dietary PUFAs could be used to modulate the lipidome of target cells to influence ferroptosis either in pathophysiological and physiological responses or in combination with therapy. However, the extent to which differences in dietary fatty acid composition might impact ferroptosis susceptibility on the major classes of immune cells is relatively unknown.

There is considerable interest in inducing ferroptosis to treat cancer (5). Importantly, recent *in vivo* preclinical studies highlight the potential of targeting specific components of the cellular pathways that limit lipid peroxidation and thereby by safeguard against ferroptosis to promote the elimination of tumours (17, 18). Despite their critical role in ferroptosis, strategies aimed at modulating cellular PUFA content *in vivo* with the goal of sensitizing cells to ferroptosis have received very little attention (10, 15). This approach is potentially very attractive as: (i) it could greatly enhance the efficacy of ferroptosis-inducing drugs, especially for cell types with low endogenous PUFA levels; (2) it could improve drug tolerability by reducing the drug concentration required for effective target cell killing; (3) supplementation with highly unsaturated PUFAs is safe (i.e. no negative effect on cell viability without GPX4 inhibition) and simple to implement, providing a very rapid route to clinical application.

Supplementing the diet with n-3 PUFAs (e.g. fish oils) has been extensively investigated for decades and, importantly, exogenous PUFAs are very effectively incorporated into cellular PLs (12, 13). Despite a wealth of studies investigating n-3 PUFA (e.g. EPA, DHA) supplementation on immune cell biology, in particular T cells, these studies have typically assessed the extent of lipidome re-remodeling in terms of changes in overall cellular fatty acid profiles. Our data add to these previous findings, demonstrating that feeding for 8 weeks with a diet enriched in EPA and DHA results in the widespread incorporation of these fatty acids into PLs from multiple classes. More importantly, by performing lipidomics on multiple immune cell types from different locations, our findings demonstrate that the incorporation of dietary EPA and DHA into immune cells is not uniform. Specifically, while B cells, monocytes and neutrophils all displayed marked (>2-fold) changes in PL-PUFA-EPA and PL-PUFA-DHA content, T cells were notably less responsive. The reasons for this are unclear but may relate to differences in the expression of fatty acid transporters or the enzymes that mediate the activation and incorporation of EPA and DHA into cellular PLs between immune cells. Consistent with our previous work (7), we show that endogenous levels of DHA vary substantially between immune cell types, typically being around 5-fold higher in lymphoid *vs* myeloid cells. Importantly, despite differences in endogenous DHA levels, exogenous DHA supplied via the diet was effectively incorporated into cellular PLs. For example, in absolute terms, B cells incorporated more exogenous DHA into PL-PUFA-DHA than other immune cells, but neutrophils and monocytes which have the lowest endogenous levels, show the largest proportional increase (∼3-fold) in PL-PUFA-DHA content in response to the High PUFA diet. These data are important as they demonstrate that despite marked differences in the regulation of endogenous cellular PL-PUFA-DHA levels between cell types, likely mediated via differences in PUFA metabolic enzyme expression, exogenous DHA supplied via the diet effectively increase PL-PUFA-DHA levels. In the context of enhancing the susceptibility of cancer cells to ferroptosis, these observations suggest that dietary provision of highly unsaturated PUFAs is likely to be effective regardless of intrinsic PUFA regulation within the cell.

A surprising outcome of the current work was that the response to the High PUFA diet differed between location. Specifically, adaptive immune cells within bone marrow and lymph nodes were more responsive to diet-induced changes than those within the blood and far more responsive than innate immune cells. Interestingly, despite marked increase in PL-PUFA-EPA and PL-PUFA-DHA levels, monocytes were not sensitized to ferroptosis by the High PUFA diet. Similarly, neutrophils, which also showed increased PL-PUFA-EPA and PL-PUFA-DHA levels following the High PUFA diet, were only susceptible to ferroptosis when found in the blood, despite the fact that lipidomic changes were greater in bone marrow neutrophils. It is possible that the resistance of bone marrow neutrophils to ferroptosis may be due to the necessity to prevent wide-spread cell death within this important organ. Interestingly, we have previously shown that the NLRP3 inflammasome is minimally activated in bone marrow neutrophils compared to blood, supporting the idea that minimizing inflammation and cell death within the bone marrow is important (19). Again, relating these findings back to enhancing the susceptibility of cancer cells to ferroptosis, this is an important observation as it indicates that the extent of PL re-modelling in response to diet is influenced by the niche in which cells reside and it cannot assumed that all cells are equally influenced by diet to the same extent. The reason why bone marrow and lymph node cells are more responsive to the diet-induced changes is unclear but may relate to the local concentration of exogenous lipids, the expression of transporters and enzymes required to incorporate the lipids, or the metabolic demands of the cells. Furthermore, after being taken up by the gut, most lipids are transported via the lymphatic system, potentially increasing the exposure of lymph node-resident immune cells to the diet.

While the focus of the current work was to determine if increasing the content of dietary EPA and DHA led to enhanced ferroptosis susceptibility, our data can equally be interpreted from the opposing perspective, i.e. can a diet that does not contain highly unsaturated PUFAs promote ferroptosis resistance. This could be relevant in conditions such as ischemia-reperfusion injury where ferroptotic cell death is implicated. Accordingly, our data support the idea that reducing or eliminating highly unsaturated PUFAs from the diet can be effective at reducing lipid peroxidation and ferroptosis, as previously demonstrated in T cells (16). With regards to future studies, it will be important to determine the effectiveness of dietary supplementation with high unsaturated PUFAs to modulate the lipid composition and, importantly, the ferroptosis susceptibility of cancer cells *in vivo*. Early evidence suggests this approach has promise (10), but, as we have shown herein, cell type and location have a considerable impact on diet-induced changes to cellular lipid content and ferroptosis susceptibly. Given the substantial and varied metabolic re-wiring that occurs in cancer cells, it would be expected that different cancer types will be differentially responsive to dietary-induced changes in lipid composition and ferroptosis susceptibility. In addition, it will be important to determine if further increasing the amount or type (e.g. AA) of highly unsaturated PUFAs in the diet can further enhance ferroptosis susceptibility. In summary, our study provides evidence for simple dietary changes that can be used to impact the lipidome of immune cells to dictate their PL-PUFA composition and render them susceptible to ferroptosis. This dietary modulation could be leveraged to enhance existing therapies or in combination with future therapies directed to induce ferroptosis across several diseases.

## Acknowledgements

We would like to thank the staff of ARA Animal Services for their exceptional care of the animals used in this study. We are extremely grateful to the following funding sources: National Health and Medical Research Council of Australia (grants GNT1189012 (to G.I.L.), GNT1194329 (to A.J.M.), GNT1197190 (to K.H.) and GNT2009965 (to P.J.M.)), the Jack Brockhoff Foundation (Bold Futures – Early Career Medical Research award #5111 (to P.K.M.)) and the Victorian Government’s Operational Support Program. The funders had no role in study design, data collection and analysis, decision to publish or preparation of the manuscript.

## Materials and methods

### Animals and diets

All procedures complied with the national guidelines for laboratory animal care and were approved by Alfred Research Alliance (ARA) Animal Ethics Committee (Melbourne, Australia). Eight-week-old male C57BL6/J mice were fed either a Control diet (AIN93G) or a High PUFA diet (SF12-037) for 8 weeks. Diets contained a fixed 16% energy from lipids, were formulated to meet known dietary requirements of omega 6 and omega 3 fatty acids, and purchased from Speciality Feeds, Glen Forrest, Australia; the fatty acid compositions of the diets are provided in **Supplementary Table 1**.

### Cell isolation and conditions

Mice were euthanised by CO_2_ asphyxiation. Blood was collected by cardiac puncture into heparin (McFarlane Med & Scientific, 13233-DB), and tibias and femurs were isolated. Bone marrow was flushed with RPMI-1640 media (GLUTAMAX) through a 40-μm filter. Red blood cells were lysed (1x RBC lysis buffer; ThermoFisher, 00-4300-54), and leukocytes were recovered following centrifugation (500g, 5mins). Total leukocyte count was obtained using an automated haematology analyser (Sysmex XS-1000i).

Cells (5 x 10^6^ per well) were seeded in 96-well plates and treated with ML210 (Sigma-Alrich, SML0521) at 10nM-20μM for 6 h or 24 h at 37°C, 5% CO_2_ in RPMI supplemented with 5% heat-inactivated Foetal Bovine Serum (FBS; Life Technologies).

### Flow cytometry

Cells were washed, pelleted (500g, 5mins) and stained in FACs buffer (HBSS without Ca^2+^/Mg^2+^, 0.1% BSA and 0.5 mM EDTA) containing anti-mouse antibodies CD45, CD11b, CD19, CD3, Ly6C and Ly6G antibodies (1:400) for 30 mins on ice in the dark. Antibody details are listed in the **Supplementary table 2**. Cells were analysed on an LSRII Fortessa (BD Biosciences) flow cytometer with FACSDiva software and processed using FlowJo.

Leukocytes were identified by FSC-A versus SSC-A, with doublets excluded by SSC-H versus SSC-A. CD45⁺ cells were subdivided into CD11b⁺ myeloid and CD11b⁻ lymphoid populations. Neutrophils were defined as CD11b⁺Ly6G⁺Ly6C⁻ and monocytes as CD11b⁺Ly6G⁻Ly6C⁺. T cells and B cells were identified as CD3⁺ and CD19⁺ within the CD11b⁻ gate. Viability was assessed using DAPI exclusion. Lipid peroxidation was measured in the FITC channel (∼510 nm) using Liperfluo.

### Lipid peroxidation and cell viability analysis

For liperfluo staining, cells were incubated with 2 μM of Liperfluo (Dojindo, L248) during the final 30 min of treatment at 37°C, washed and surface-stained prior to flow cytometric analysis. For viability assessment, cells were resuspended in FACS buffer containing 0.05 μg/mL DAPI solution (BD Pharmingen, 564907) and incubated for 5 mins at room temperature immediately before acquisition; DAPI (-) cells were considered viable cells.

### Fluorescence-activated cell sorting for lipidomics

Leukocyte subsets were sorted using a BD FACSAria, FACSAria Fusion or Influx at the ARA Flow Cytometry Core Facility. Flow antibodies and gating strategies are detailed in **Supplementary Table 2**. For sorting, gating strategies were first set up based on FSC-A versus SSC-A, SSC-H versus SSC-A (doublet exclusion) and side scatter area versus viability dye (viable cell isolation). Sorted event thresholds were set to 5 × 10⁵ cells for neutrophils, B cells and monocytes, and 1–1.5 × 10⁵ cells for T cells. Sorted cells were washed in PBS (without Ca²⁺/Mg²⁺) and stored at −80 °C.

Cells were sonicated, protein quantified, and lipids extracted using a single-phase butanol/methanol method as described previously¹. Plasma QC samples were incorporated every 10 samples, and pooled technical QC samples every 20 injections. Lipid species were analysed using an Agilent 6495C triple quadrupole mass spectrometer coupled to an Agilent 1290 HPLC with a C18 column (45 °C), using scheduled multiple reaction monitoring in positive and negative ion modes. A total of 777 lipid species were quantified relative to internal standards (Supplementary Table 5). Chromatographic and mass spectrometry parameters are detailed in Supplementary Tables 3–4.

### Lipid extraction

Sorted cells were thawed and sonicated using an S-400 sonicator (Misonix) at an amplitude of 13% for 25 seconds. Protein concentration was determined by Quant-iT Protein Assay Kit (ThermoFisher Scientific) as per manufacturer’s protocol. To prepare for lipid extraction, the cell homogenates were stored at -80°C for 2 hours and dried down overnight in a SpeedVac (Thermo Fisher Scientific).

Lipids were extracted using a single-phase butanol/methanol extraction method as described previously (7, 20). Briefly, in addition to sample lipid extracts, after every 10 samples, 1 plasma QC (PQC) sample consisting of a pooled set of plasma samples taken from 6 healthy individuals and extracted alongside the study samples were incorporated to provide a measure of technical variation. 50 *μ*L of 1:1 butanol:methanol containing internal standard mixtures were added to each sample. The samples were vortexed, sonicated in a water bath sonicator for 1 hour and centrifuged for 10 min at 13,000 x g. The supernatant was transferred into 0.2 ml micro-inserts in glass vials and stored at -80°C. A total of 5*μ*L was injected for each cell sample.

### Liquid chromatography tandem based mass spectrometry

Lipid analysis was performed as described previously with modifications(20). Analysis of lipid extracts was performed on an Agilent 6495C QQQ mass spectrometer with an Agilent 1290 series HPLC system and a single ZORBAX eclipse plus C18 column (2.1x100mm 1.8mm, Agilent) with the thermostat set at 45°C. The running solvent consisted of solvents A and B containing of water: acetonitrile: isopropanol in the ratios of 5:3:2 and 1:9:90 respectively, including 10 mM ammonium formate (in solvents A and B) and 5μM medronic acid (in solvent A only). The chromatography and mass spectrometer settings are detailed in **Supplementary Table 3** and **Supplementary Table 4**. In addition to sample extract and PQC, technical QC samples (TQC) consisting of PQC extracts which were pooled, then divided into individual vials was added to the run (at a ratio of 1 TQC per 20 samples) to provide a measure of technical variation from the mass spectrometer only. Relative quantification of lipid species was determined by comparison to the relevant internal standard in **Supplementary Table 5**. Acyl annotations were derived directly from fragmentation for PE plasmalogens (PE P), while other glycerophospholipids were annotated through elution time matching from using alternative ionisation methods and adducts as previously described (7, 20).

### Statistical analysis

Flow cytometry and lipidomic data were analysed using one- or two-way ANOVA with Tukey’s post hoc correction (GraphPad Prism or R) or unpaired Student’s *t* tests. Analysis of lipidomic data was performed using R version 4.4.1 (CRAN). Uniform Manifold Approximation and Projection (UMAP, ‘umap’ package (v0.2.10.0) in R (v4.4.1)) was applied to log10-transformed, mean centered and standard deviation scaled lipid concentration data for visualization of the global lipidomic profiles of immune cells between dietary conditions at different tissue sites. Differences in the concentrations of lipid classes and individual lipid species between groups were assessed using unpaired Student’s t tests performed on log10-transformed data. Resulting P values were adjusted for multiple comparisons using the Benjamini-Hochberg method. Statistical significance was determined using a false discovery rate (FDR) threshold of 10%. Lipid species that were significantly different between dietary conditions in at least one immune cell type were included in the heatmap visualization. For forest plots, differences in lipid concentrations between groups were assessed using unpaired Student’s *t* tests performed on log_10_-transformed data. Resulting *P* values were adjusted for multiple comparisons using the Benjamini-Hochberg False Discovery Rate correction with a threshold of 0.1.

---

**Fig. S1.**
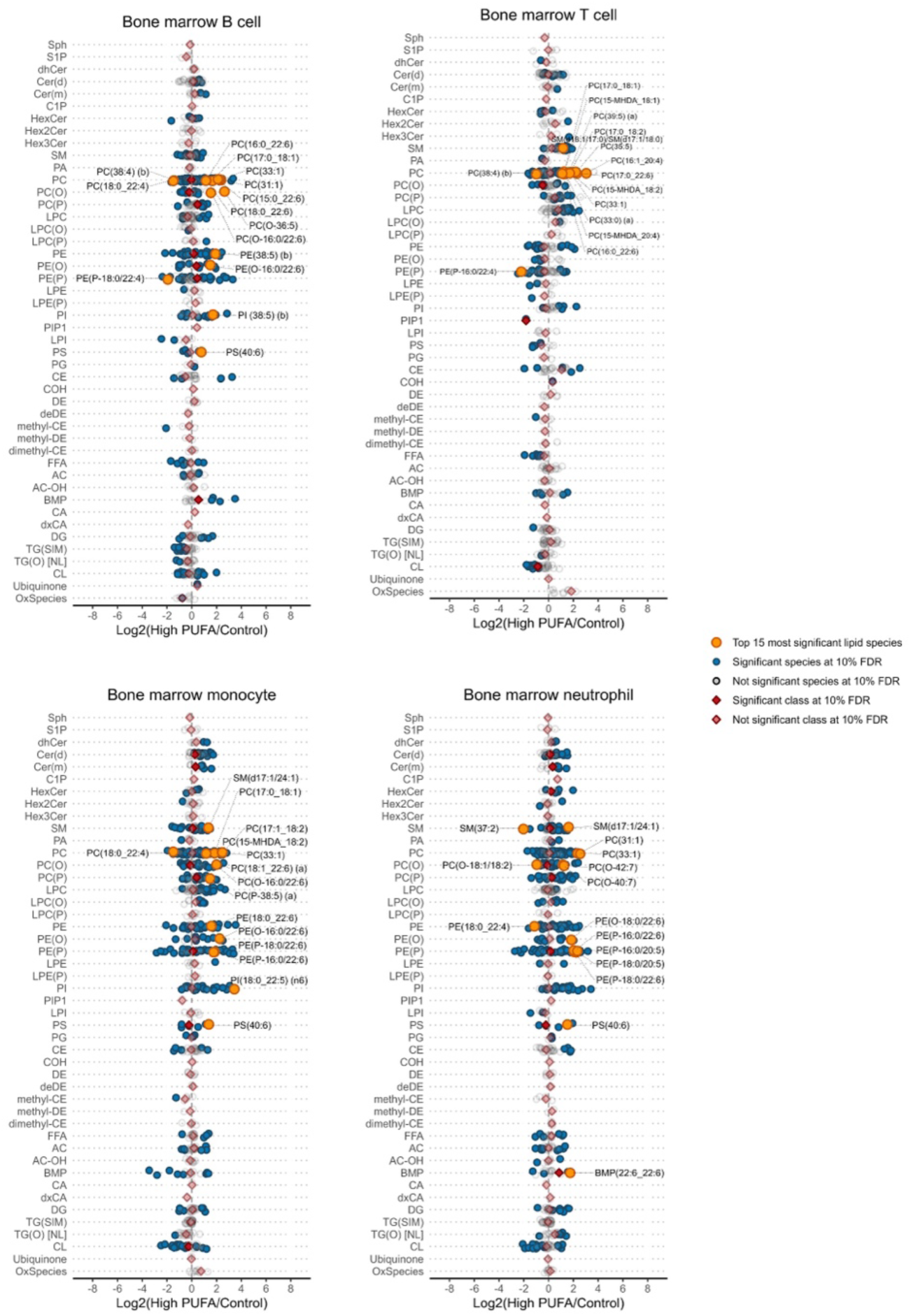
A High PUFA diet alters bone marrow immune cell lipid composition. Forest plots of individual lipid species and class totals in immune cells from the bone marrow (741 measured lipid species) of mice fed either a Control diet or High PUFA diet for 8 weeks. The following number of lipids were significantly different: B cells, 274; T cells, 221; monocytes, 343; neutrophils, 333. n = 7-8 mice per group. Data are presented as fold change and error bars are 95% confidence intervals.

**Fig. S2.**
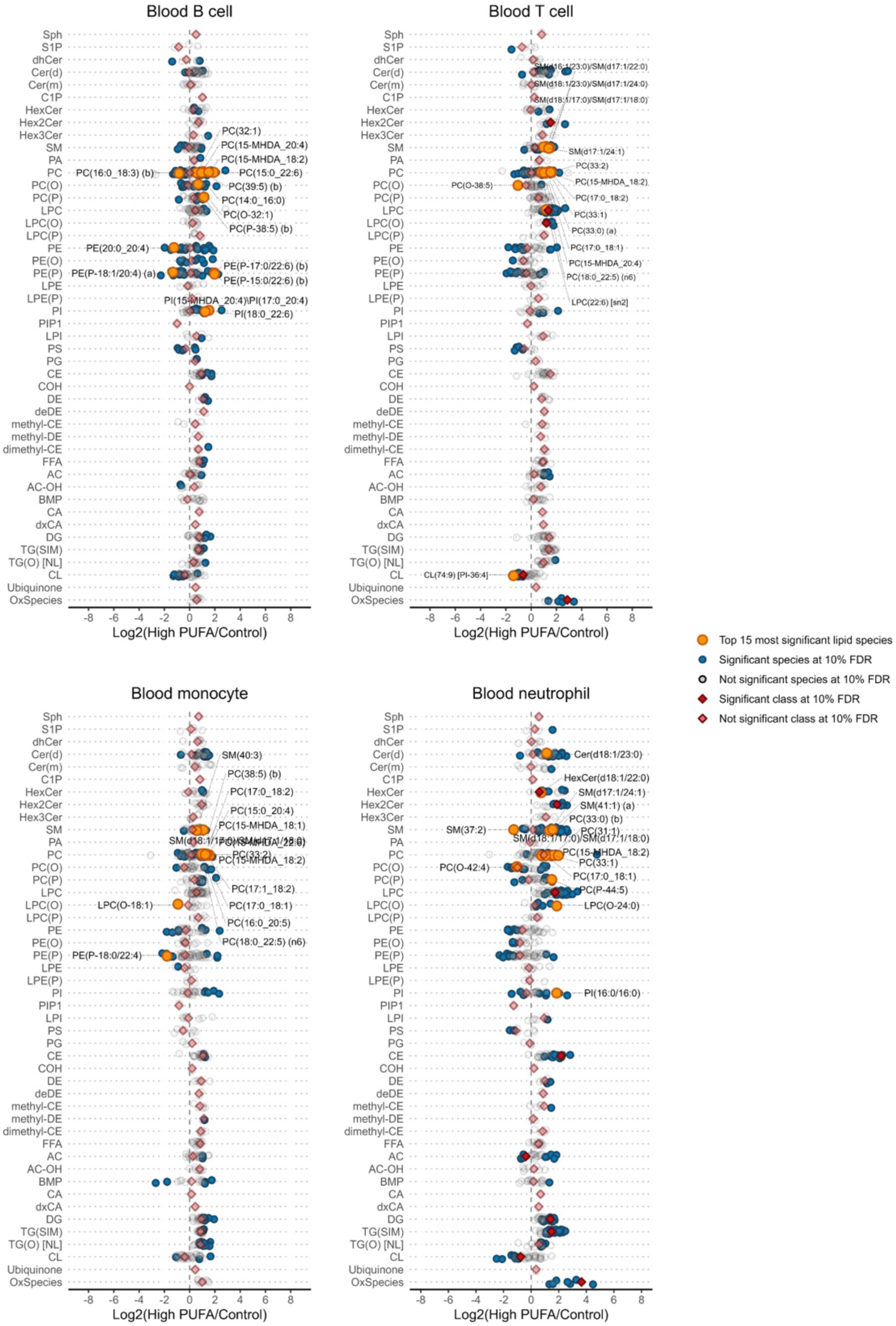
A High PUFA diet alters blood immune cell lipid composition. Forest plots of individual lipid species and class totals in immune cells from the blood (741 measured lipid species) of mice fed either a Control diet or High PUFA diet for 8 weeks. The following number of lipids were significantly different: B cells, 196; T cells, 154; monocytes, 129; neutrophils, 257. n = 6-8 mice per group. Data are presented as fold change and error bars are 95% confidence intervals.

**Fig. S3.**
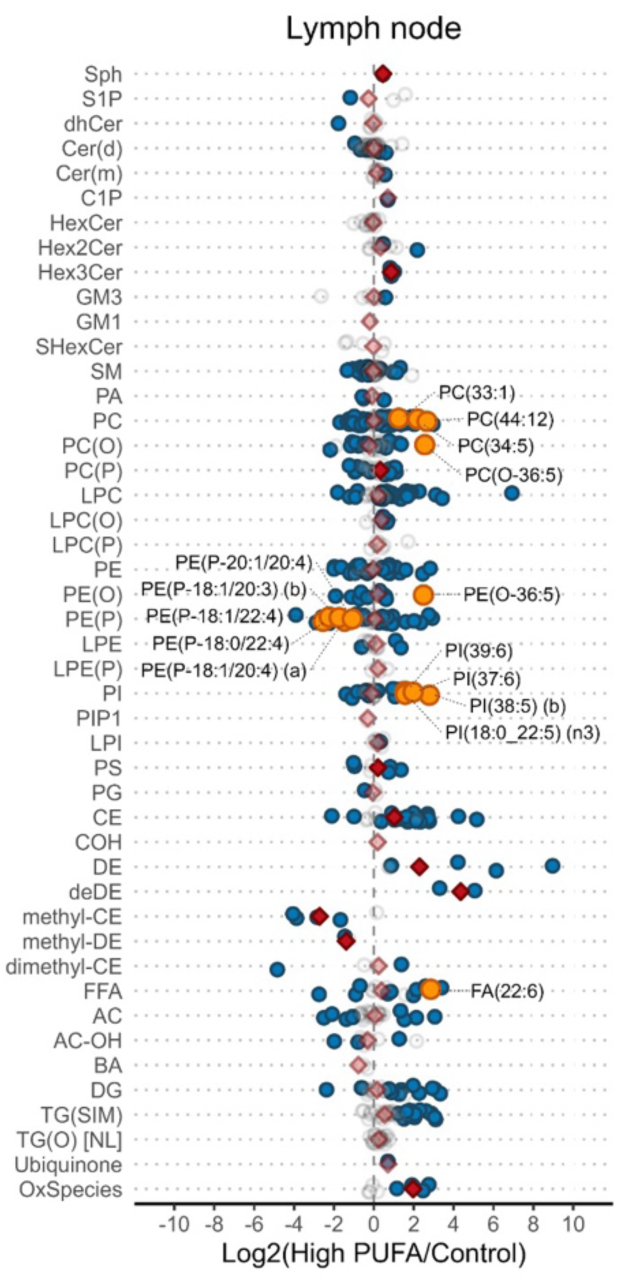
A High PUFA diet alters lymph node immune cell lipid composition. Forest plots of individual lipid species and class totals in immune cells from the lymph nodes (714 measured lipid species) of mice fed either a Control diet or High PUFA diet for 8 weeks. 360 lipids were significantly different. n = 8 mice per group. Data are presented as fold change and error bars are 95% confidence intervals.

**Fig S4.**
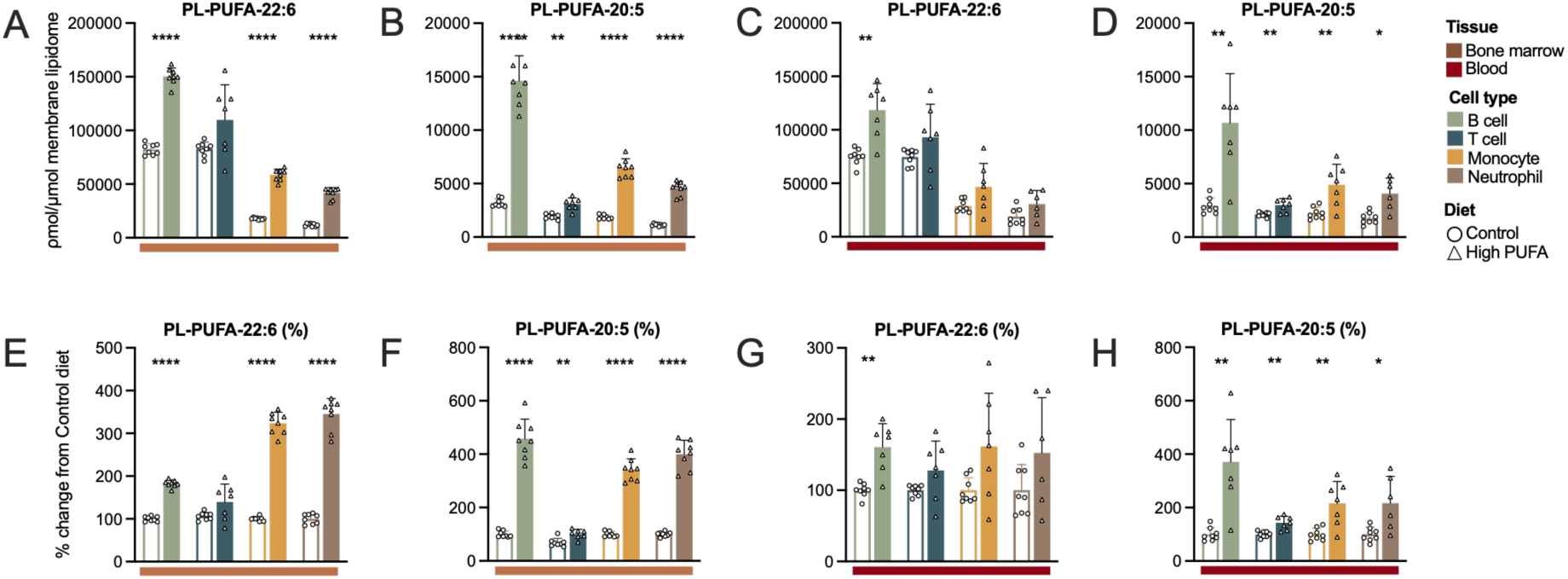
A High PUFA diet increases the overall content of PLs containing 20:5 and 22:6. **A-D** Concentration and **E-H** relative percentage change in overall levels of PLs containing 20:5 and 22:6 PUFAs in **A, B, E, F** B cells, T cells, monocytes and neutrophils from the bone marrow and **C, D, G, H** blood of mice fed either a Control diet or High PUFA diet. An independent samples t-test was used to compare cell types between diets. *p<0.05, **p<0.01, ***p<0.001 and ****p <0.0001. n = 7-8 mice per group for bone marrow immune cells and n = 6-8 for blood immune cells mice per group.

**Fig. S5.**
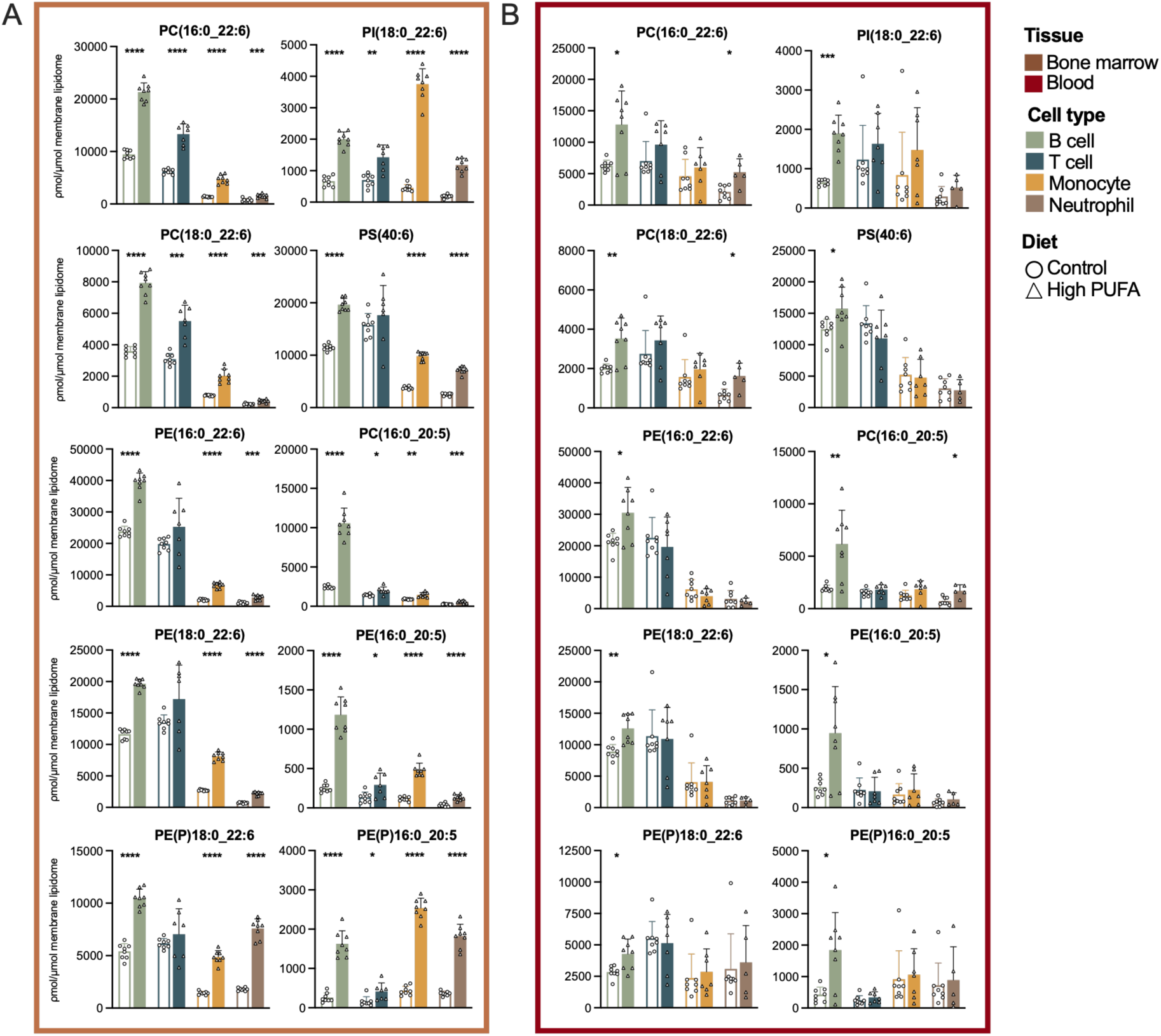
A High PUFA diet increases the abundance of PUFA-containing PL species from multiple PL classes in immune cells. **A, B** Concentrations of the indicated PUFA-containing phospholipid species determined by MS lipidomics in B cells, T cells, monocytes and neutrophils from the **A** bone marrow and **B** blood of mice fed either a Control diet or High PUFA diet. An independent samples t-test was used to compare cell types between diets. *p<0.05, **p<0.01, ***p<0.001 and ****p <0.0001. n = 7-8 mice per group for bone marrow immune cells and n = 6-8 mice per group for blood immune cells.

